# Adaptation modulates correlated response variability in visual cortex

**DOI:** 10.1101/087049

**Authors:** Nathaniel C. Wright, Mahmood S. Hoseini, Ralf Wessel

## Abstract

Cortical sensory responses are highly variable across stimulus presentations. This variability can be correlated across neurons (due to some combination of dense intracortical connectivity, cortical activity level, and cortical state), with fundamental implications for population coding. Yet the interpretation of correlated response variability (or “noise correlation”) has remained fraught with difficulty, in part because of the restriction to extracellular neuronal spike recordings. Here, we measured response variability and its correlation at the most microscopic level of electrical neural activity, the membrane potential, by obtaining dual whole-cell recordings from pairs of cortical pyramidal neurons during visual processing. We found that during visual stimulation, correlated variability adapts towards an intermediate level and that this correlation dynamic is mediated by intracortical mechanisms. A model network with external inputs, synaptic depression, and structure reproduced the observed dynamics of correlated variability. These results establish that intracortical adaptation self-organizes cortical circuits towards a balanced regime at which network coordination maintains an intermediate level.

---

Sensory cortex is not simply one layer in a feedforward network (Fig. 1a); it receives strong inputs from thalamus, but intracortical feedback dominates cortical circuitry (Fig. 1b). This tangle of cortical connections causes neural activity to be coordinated across multiple spatial and temporal scales^1,2^. Moreover, in a given cortical network, the strength of this coordination can vary with activity level and network state^3–8^, which is considered to have implications for cortical function^9–12^. For example, weak coordination corresponds to a larger “library” of words available to the spatiotemporal code, while stronger coordination supports signal propagation (Fig. 1c). The realized level of coordination in active cortical circuits is expected to represent a balance between such competing system needs. Two unanswered questions concerning coordination continue to block our path to understanding sensory processing in cerebral cortex. First, what are the levels of cortical coordination during sensory processing, and to what extent do these levels change with varying stimulus conditions (Fig. 1c)? Second, what mechanisms are responsible for the realized level of cortical coordination and its changes?

**Figure 1.**
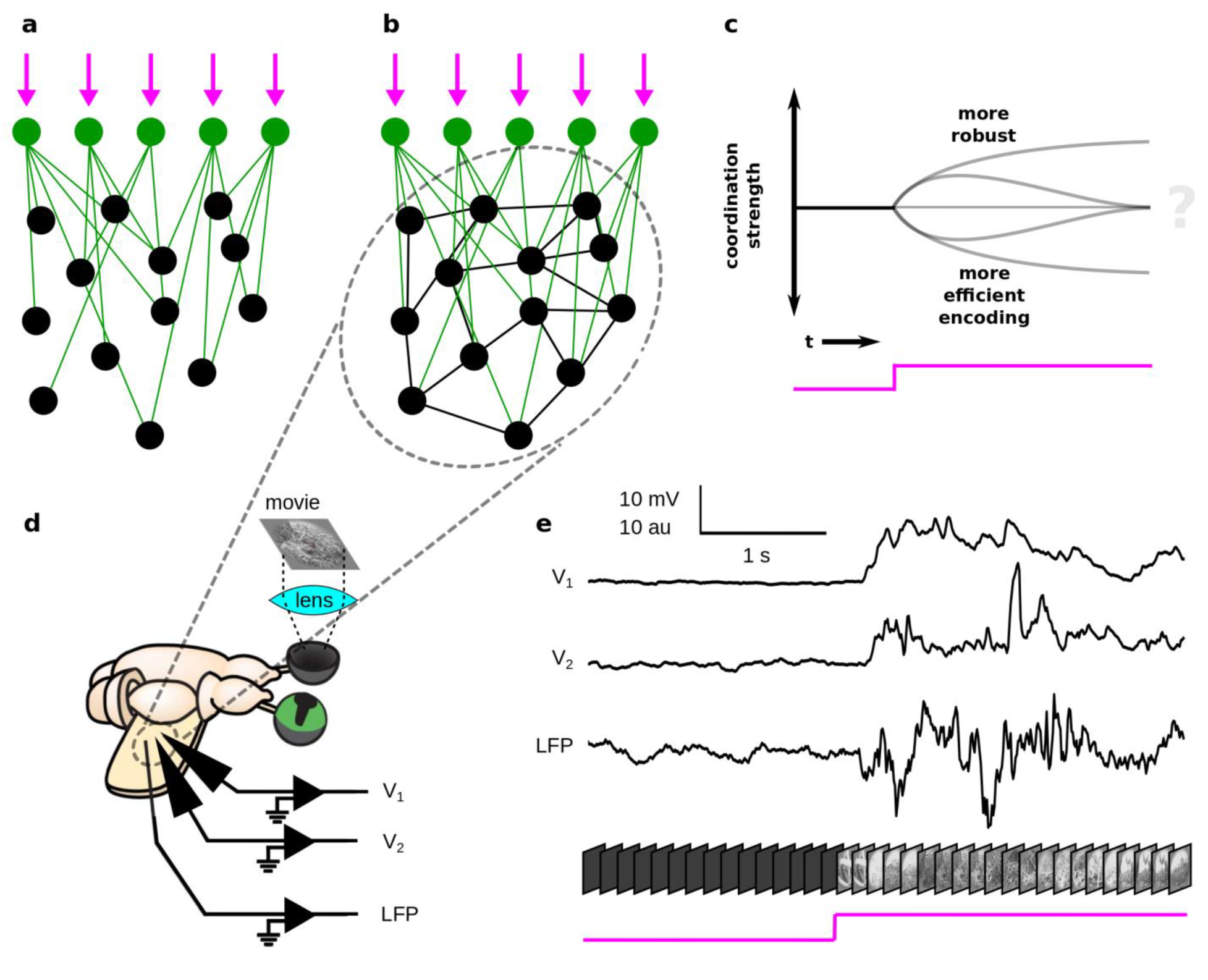
Investigating the dynamics of correlated variability in recurrent circuits of visual cortex. (a) Feedforward thalamocortical network subject to sensory inputs (magenta). Coordination between pairs of cortical neurons (black) is determined by convergence patterns in thalamic inputs (green). (b) A more realistic, interaction-dominated thalamocortical network, in which the inputs to any one cortical neuron arise primarily from other cortical neurons. Coordination is thus a function of both feedforward and recurrent inputs. (c) The level of cortical coordination affects cortical function, and it is unknown if and how this changes with sensory stimulation. (d) We simultaneously recorded the membrane potentials from pairs of cells, as well as the nearby LFP, during ongoing and visually-evoked activity in a densely-interconnected thalamocortical network. (e) Pairwise membrane potential recordings provided a measure of cortical coordination across stimulus conditions that avoids the pitfalls of spike data.

Recording the spike trains from pairs of neurons for repeated presentations of an identical stimulus and evaluating the level of correlated response variability (or “noise correlation”) has remained the work horse of investigating cortical coordination (see Cohen, Kohn 2011, and Doiron, Kumar 2016 for reviews). This spike-based approach is popular in part because of the relative ease of obtaining spiking responses from pairs of neurons in intact brains. Reported values of spike-based noise correlation tend to be significantly nonzero, but results have varied across studies^4,6,13–15^. Furthermore, the interpretation of cortical coordination from spike data is littered with complications, including a spike-rate dependence of noise correlation values^15,16^, the underrepresentation of sparse-spiking neurons, and possible biases introduced by the spike-sorting process^15,17^. In conclusion, the important study of cortical coordination, including its relation to mechanisms and function, has been restricted by its focus on spike recordings, and continues to represent an unmet challenge in systems neuroscience.

In response to this need, we investigated the dynamics of correlated response variability at the level of the membrane potential by obtaining dual whole-cell recordings from pairs of cortical pyramidal neurons during visual processing (Fig. 1d, e). We found a high level of trial-to-trial membrane potential response variability. Further, correlated variability in the gamma band range of membrane potential fluctuations increased at stimulus onset, but returned to pre-stimulus values during continued visual stimulation. A brief visual stimulus, triggering persistent cortical activity, elicited a similar dynamic of correlated variability, thus implicating an intracortical mechanism. A model network with small-world connectivity, external inputs, and synaptic depression reproduced the observed dynamics of correlated variability, and further indicated that network oscillations play a crucial role. Taken together, these results establish adaptation towards an intermediate level of coordination as a fundamental principle of cortical organization during visual processing.

## RESULTS

To quantify response variability and its correlation across neurons, we recorded the membrane potential (V) from 35 pairs of pyramidal neurons in visual cortex of the turtle *ex vivo* eye-attached whole-brain preparation during visual stimulation of the retina (Fig. 1d). Ongoing activity in turtle visual cortex was largely quiet. In contrast, visual stimuli evoked barrages of postsynaptic potentials in cortical pyramidal neurons that were accompanied by extensive fluctuations in the nearby local field potential recording (Fig.1e), indicating strong concurrent network activity.

### Pyramidal neuron membrane potential visual responses are highly variable

We recorded from 19 pairs of pyramidal neurons while presenting continuous visual stimulation, and from 16 pairs while presenting brief flashes (see Methods). Single-neuron membrane potential responses to repeated presentations of extended stimuli varied from trial to trial, with a response variability magnitude that exceeded the trial-averaged mean response (Fig. 2a). Importantly, the magnitude of the response variability was qualitatively unchanged when the visual stimulus consisted of brief flashes, which evoked long-lasting responses in visual cortex (Fig. 2b). This stimulus invariance of the trial-to-trial response variability indicates an intra-thalamocortical origin of the network activity and the resulting membrane potential fluctuations.

**Figure 2.**
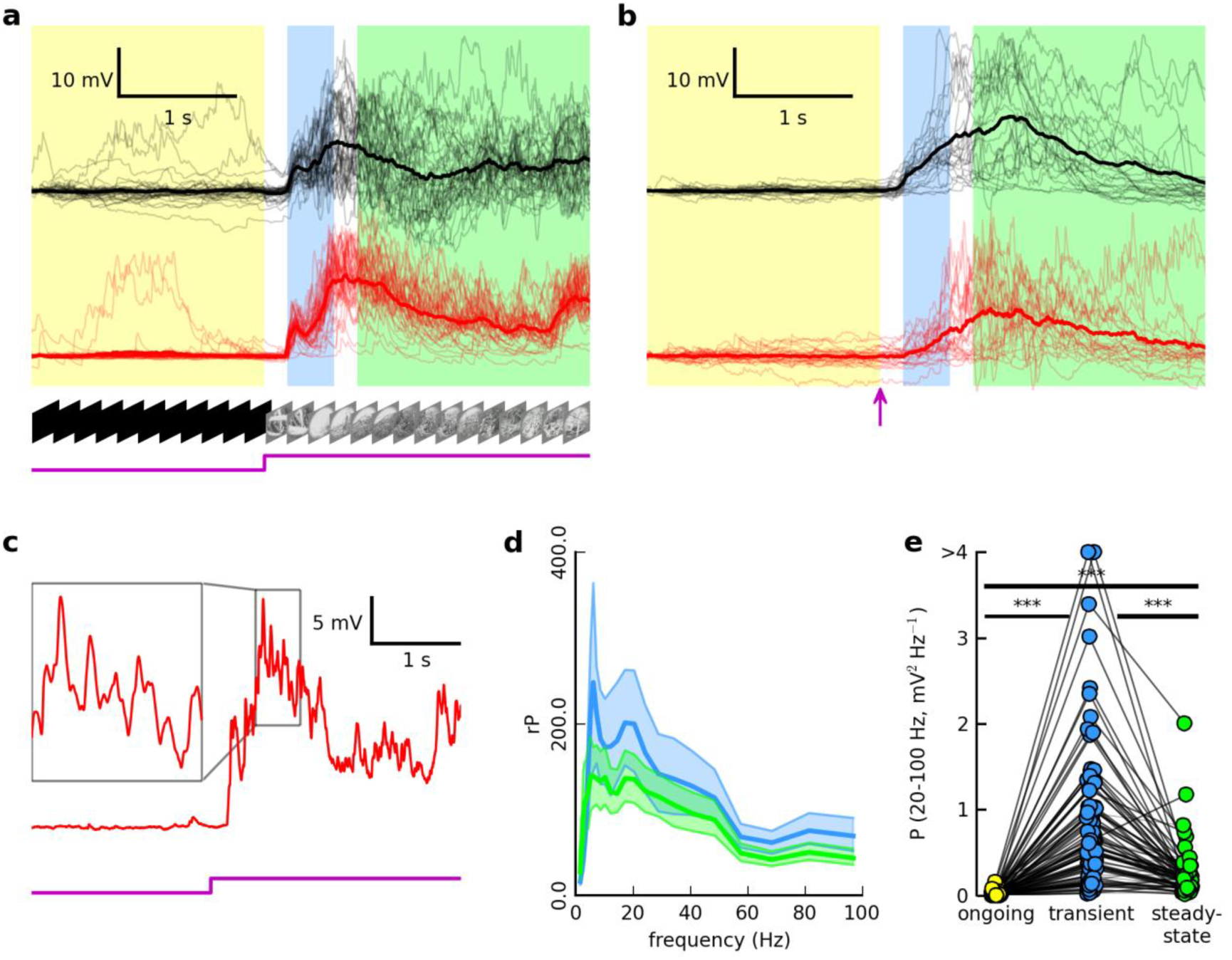
Dynamics and complexity of trial-to-trial response variability. (a) Single-trial responses (low opacity) and across-trial average responses (high opacity) for two simultaneously-recorded neurons. Stimulus is naturalistic movie (see Methods). Single trials artificially aligned for clarity. (b) Same as in (a), but for a different pair of cells, and stimulus is 150 ms red (640 nm) whole-field flash, with onset at magenta arrow, (see Methods). (c) Single-trial response from cell in (a). Inset: high-frequency activity nested within the broader depolarization. (d) Average relative power spectrum (evoked power divided by ongoing) of residuals for red traces in (a) for the transient (blue) and steady-state (green) epochs. Shaded regions indicate +/− 95% confidence intervals by bootstrapping method. (e) Trial-averaged gamma (20 – 100 Hz) power for 79 cells, for brief and extended visual stimulation. Each dot represents the across-trial average gamma power for one cell for that epoch. Dark (light) lines connecting dots represent (in)significant changes in gamma power for that cell across epochs. Across-epoch significance for each cell determined by comparing bootstrap intervals. Asterisks above line connecting two epochs indicates results of Wilcoxon signed-rank significance test for difference in populations of values for those epochs (**: 0.001 ≤ P < 0.01; ***: P < 0.001; no asterisk and gray line: P > 0.05). Two outliers truncated for clarity.

For any given trial of visual stimulation, the evoked membrane potential fluctuations were large and consisted of high-frequency fluctuations nested within broader deflections (Fig. 2c). To quantify the frequency content of the single-trial fluctuations from the mean response, we first calculated the membrane potential residual (V_r_), which is the single-trial membrane potential recording from which the trial-averaged membrane potential time series has been subtracted. We then divided the evoked residuals into two analysis windows: the transient (200 to 600 ms after stimulus onset) and steady-state (800 to 2800 ms after stimulus onset) windows (**Supplementary Information 1**). Finally, we calculated the relative power spectral density (rP), which is the power spectral density of the membrane potential residual for the transient or steady-state window divided by its trial-averaged counterpart from the ongoing window (2 s prior to stimulus onset).

This analysis revealed four important features concerning the spectral content of the residual membrane potential fluctuations and of the trial-to-trial response variability. First, evoked power of residual membrane potential fluctuations in the 0.1 to 100 Hz range typically increased by two orders of magnitude compared to ongoing activity (Fig. 2d, **Supplementary Fig. 2**). Second, the frequency content of the membrane potential residual varied across trials (as indicated by the broad confidence bands in Fig. 2d). Third, the relative power spectral density typically consisted of a prominent peak located approximately in the 4 to 10 Hz theta range and a broader, but distinct, distribution in the 20 to 100 Hz gamma range. Fourth, for both movies and flashes, gamma power increased from the ongoing to the transient window, and significantly decreased from transient to steady-state (Fig. 2e). Gamma power of the transient activity varied drastically across cells.

Together, these data establish that cortical pyramidal neuron membrane potential visual responses (i) have complex temporal dynamics, (ii) are highly variable from trial to trial, and (iii) differ from neuron to neuron (Fig. 2).

### Correlated variability adapts during visual stimulation

The complex and extensive variability of membrane potential visual responses (Fig. 2) and the interconnected nature of cortical circuits (Fig. 1b) raised the question to what extent the response variability is correlated across pyramidal neurons. To address this question, we calculated the Pearson correlation coefficient between residual membrane potential fluctuations for each trial and window of interest, i.e., the ongoing, transient, and steady-state windows. We focused on gamma-band (20 – 100 Hz) activity, which captures the fast, nested membrane potential fluctuations (Fig. 2c, inset, Fig. 3a, and see **Supplementary Information 2**). This band of activity is thought to be associated with narrow “windows of opportunity” for spiking, determining the precise timing of spikes within a broader depolarization (see Haider, McCormick, 2009 for a review). Trial-averaged correlation coefficients (CC) for ongoing activity were broadly distributed across pairs of pyramidal neurons (Fig. 3b), and the population average (<CC>) was significantly nonzero (<CC> = 0.03, P = 0.006, one-sided t-test). In response to continuous visual stimulation, trial-averaged correlation coefficients increased significantly compared to ongoing values (Fig. 3b), to an elevated population average of <CC> = 0.11 (P = 2.9 x 10^−4^ for ongoing – transient comparison, Wilcoxon signed-rank test). In the steady-state period, i.e., during continued stimulus presentation, trial-averaged correlation coefficients returned to near-ongoing values (<CC> = 0.041, P = 1.6 x 10^−4^ for transient – steady-state comparison, P > 0.05 for ongoing – steady-state comparison).

**Figure 3.**
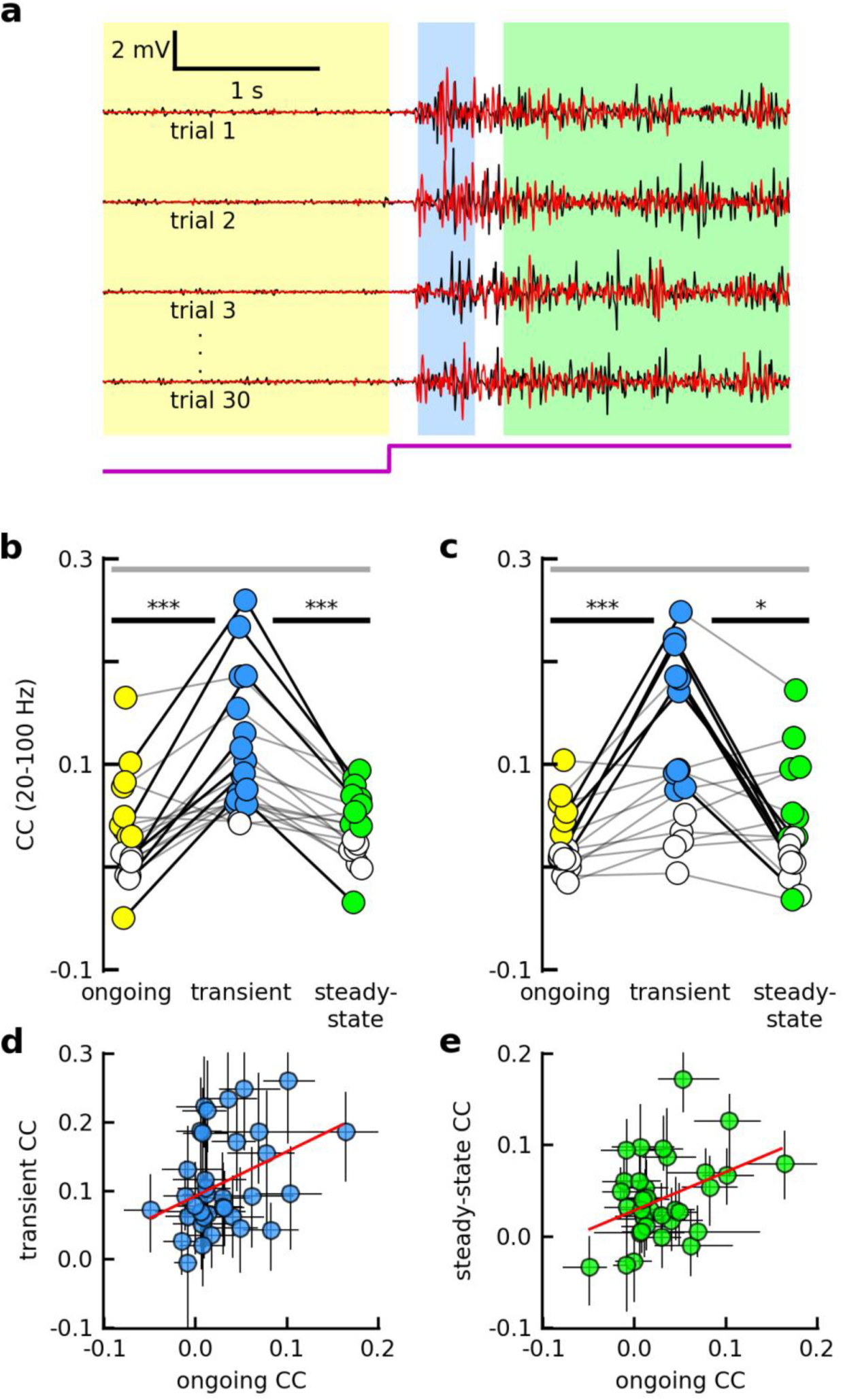
Evoked gamma band correlated variability appeared to be modulated by internal mechanisms. (a) Examples of gamma band (20 – 100 Hz) residual membrane potential pairs for several trials (same pair as in 2a). (b) Trial-averaged CC values for each of 19 pairs, 20 – 100 Hz, continuous visual stimulation (see Methods). Each dot represents the across-trial average CC value for one pair for that epoch. Colored (white) dots represent values (not) significantly different from zero (one-sided t-test). Otherwise, same as in 2e. (c) Same as in (b), but for 16 pairs, and brief visual stimulation (see Methods). (d) Across-trial average transient CC vs. average ongoing CC for each pair, for all stimuli. Error bars indicate 95% confidence intervals by bootstrapping method. Red line indicates significant linear regression fit (r^2^ = 0.14, P = 0.026). (e) Same as (d), but for steady-state epoch (r^2^ = 0.15, P = 0.02).

These results were largely robust with respect to choices of window sizes and gaps between windows (see **Supplementary Information 1** and **Supplementary Fig. 1**). These changes in correlated variability primarily reflected changes in phase synchrony in the gamma band residual activity of simultaneously-recorded neurons (see **Supplementary Information 3** and **Supplementary Fig. 4**). In contrast, low-frequency (0.1 – 20 Hz) CC values followed a different dynamic (**Supplementary Fig. 3a, b**) and were not significantly related to gamma-band CC (see **Supplementary Information 2** and **Supplementary Fig. 3c - e**).

The observed dynamics of gamma band correlated variability in response to continuous visual stimulation could be imposed by the spatiotemporal structure of the stimulus, or alternatively, could be intrinsic to the thalamocortical system. To distinguish between these two hypotheses, we recorded from 16 pyramidal neuron pairs while presenting brief flashes (1 ms – 200 ms) of light, which evoked responses lasting several seconds in the visual cortex (Fig. 2b). We found that across the population of all pairs, CC values for responses to brief stimuli were not significantly different from those for continuous stimuli (p > 0.05, Wilcoxon rank-sum test, for all epochs, see **Supplementary Information 4**, and **Supplementary Fig. 5**). Importantly, the same dynamics of correlated variability were observed for brief stimuli (<CC> = 0.03, P = 0.003 ongoing, <CC> = 0.11, P = 3.0 x 10^−5^ transient, <CC> = 0.041, P = 0.006 steady-state, one-sided t-test; P = 6.4 x 10^−4^ for ongoing – transient comparison, P = 0.020 transient – steady-state comparison, P > 0.05 ongoing – steady-state comparison, Wilcoxon signed-rank test, Fig. 3c). The similarity of the dynamics of correlated variability for brief and continuous stimuli implicates a mechanism that is stimulus-invariant and likely to be intracortical in origin.

The evidence of an intracortical origin of correlated variability suggested to us that the correlated variability of ongoing activity for a given pair should be predictive of its correlated variability in response to visual stimulation. Indeed, we found that CC values during the ongoing epoch were significantly related to CC values for both evoked epochs (r = 0.38, P = 0.03, transient vs. ongoing, Fig. 3d; r = 0.39, P = 0.02, steady-state vs. ongoing, Fig. 3e, Pearson correlation). This observation suggests a close link between the underlying mechanisms that determine ongoing and evoked correlated variability, with connectivity being one candidate mechanism^18,19^.

### Correlated variability is related to the network state

Recent experimental and computational work suggests that the level of coordination between pairs of neurons might be shaped not only by anatomical connectivity, but also by network activity level and network state^3–8^. We therefore asked how well network activity level and/or network state could explain the observed dynamics of correlated variability.

As a proxy for network activity level, we inferred the level of presynaptic spiking activity using the average gamma power in residual membrane potentials. As a population, the gamma power dynamic (Fig. 2e) was qualitatively similar to that of CC (Fig. 3b, c). In contrast, for a given pair of cells, the geometric mean of the absolute change in power (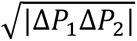) was not related to the magnitude of the change in CC amplitude for the ongoing – to – transient transition (r = 0.28, P = 0.10, Pearson correlation). The two quantities were however related for the transient – to – steady-state transition (r = 0.40, P = 0.017, Pearson correlation) (Fig. 4b). In conclusion, changes in network activity levels alone did not fully explain the changes in gamma band correlated variability.

**Figure 4.**
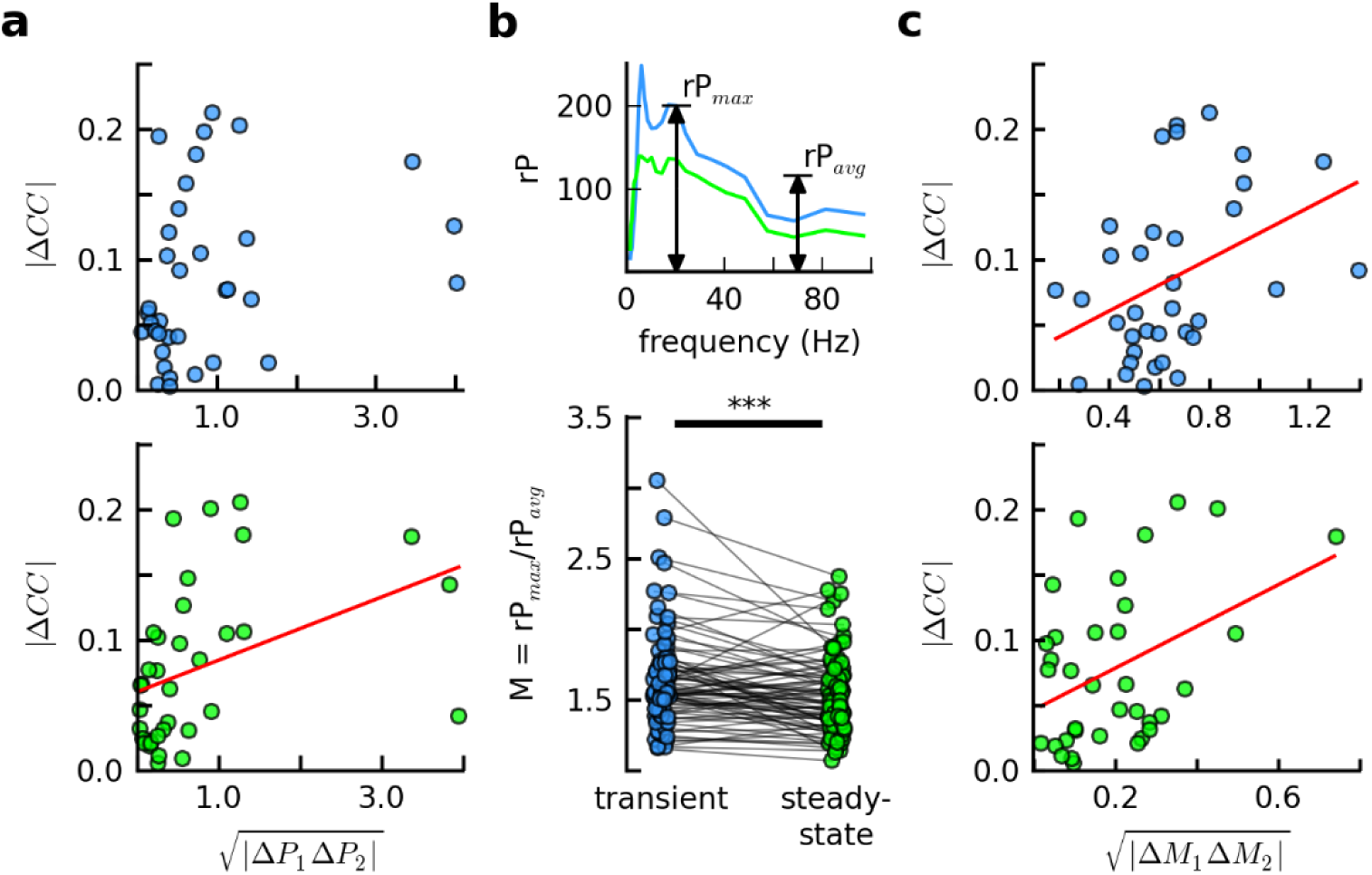
Changes in CC are related to changes in the shape of relative power spectra. (a) Absolute value of change in CC (evoked minus ongoing) vs. absolute value of geometric mean of change in gamma power, for each pair, for transient (top), and steady-state (bottom) epochs. (b) Top: same as in 2(d), but confidence bands omitted for clarity. For each cell, the peak power index (M), is defined as the maximum relative power value in the gamma range (rP_max_), divided by the average over the gamma range (rP_avg_). Bottom: M for all recorded cells (brief and continuous stimulation), for transient (blue) and steady-state (green) epochs. (c) Same as in (a), but for absolute value of geometric mean of change in M, for all cells. If present, red lines indicate significant linear regression fit (P < 0.05).

Next, we sought a measure of the network state. Specifically, we focused on the degree of action potential synchrony as an indicator of network state. Computational work has shown that synchronous network spiking can generate prominent subthreshold membrane potential fluctuations in a narrow frequency band ^20^. Here, we used a measure of power spectrum “peakiness” in the 20 to 100 Hz range as a proxy for synchronous network spiking. The residual relative power spectra (rP) of some cells tended to contain peaks in the gamma band during the transient epoch, which were often smaller or absent in the steady-state (Fig. 2d, **Supplementary Fig. 2**). To quantify this aspect of the power spectrum for each cell and epoch, we defined the quantity M as the ratio of the maximum relative power in the gamma range (rP_max_) to the average relative gamma power (rP_avg_) (Fig. 4b, **top**). Across the population of cells, M varied continuously, and the population average (<M>) decreased significantly from transient to steady-state (<M> = 1.65 transient, <M> = 1.54 steady-state, P = 2.2 x 10^−4^ for transient – steady-state comparison) (Fig. 4b, **bottom**). For the population of pairs, we found that the geometric mean of the absolute change in M (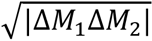) was significantly related to the amplitude of the change in CC (r = 0.40, P = 0.02 ongoing – to – transient; r = 0.41, P = 0.01 transient – to – steady-state, Pearson correlation, where Δ*M* = *M*_*trans*_ – 1 for the ongoing – transient transition) (Fig. 4c). In conclusion, for a given pair of cells, the change in network state, measured as a change in gamma spectrum “peakiness”, was an indicator of the change in correlated variability.

### Synaptic time constants, synaptic depression, and synaptic clustering together mediate the dynamics of correlated variability

What biophysical mechanisms could mediate the experimentally-observed response properties (i.e., across-trial variability (Fig. 2a, b), subthreshold gamma oscillations (Fig. 2c, 3a), the dynamics of correlated variability (Fig. 3) and its network state dependence (Fig. 4))? To address this question, we investigated a model network (Methods) of 800 excitatory and 200 inhibitory leaky integrate-and-fire neurons, with Poisson process external inputs to all excitatory neurons (Fig. 5a). Excitatory-to-excitatory and excitatory-to-inhibitory connections had small-world connectivity, with 5% connection probability. Inhibitory-to-inhibitory and inhibitory-to-excitatory synapses were random, with 20% connection probability. An increase in the external input rate mimicked the stimulus. We selected a set of synaptic rise and decay times^20,21^ that were consistent with gamma oscillations in the instantaneous network firing rate when the network was subject to strong external drive. Motivated by previous experiments^22^ and models^23^, we implemented adaptation via short-term synaptic depression with recovery (Methods). Armed with this battery of mechanisms and structural complexity, we simulated the network spiking in response to repeated stimulus presentations (20 trials).

The model network reproduced the experimentally-observed response variability and gamma oscillations. The simulated membrane potentials from randomly-selected excitatory model neurons revealed fluctuations and across-trial response variability (Fig. 5b) that were qualitatively similar to what we observed from recorded membrane potentials of pyramidal neurons (Fig. 2a, b). Furthermore, the gamma band residuals of simulated membrane potentials (Fig. 5c) largely resembled the experimentally-observed residual subthreshold gamma oscillations (Fig. 3a). Residual power spectra were peaked in the gamma range (**Supplementary Fig. 6a, top**), at frequencies that coincided with peaks in the spectra of the instantaneous network firing rate (**Supplementary Fig. 6a, bottom**). Due to synaptic depression with slow recovery, the model also reproduced the eventual decrease in the total gamma power in membrane potential residuals (**Supplementary Fig. 6b**), and in the amplitudes of gamma-band peaks in the residual membrane potential and population spike rate spectra (**Supplementary Fig. 6a**).

**Figure 5.**
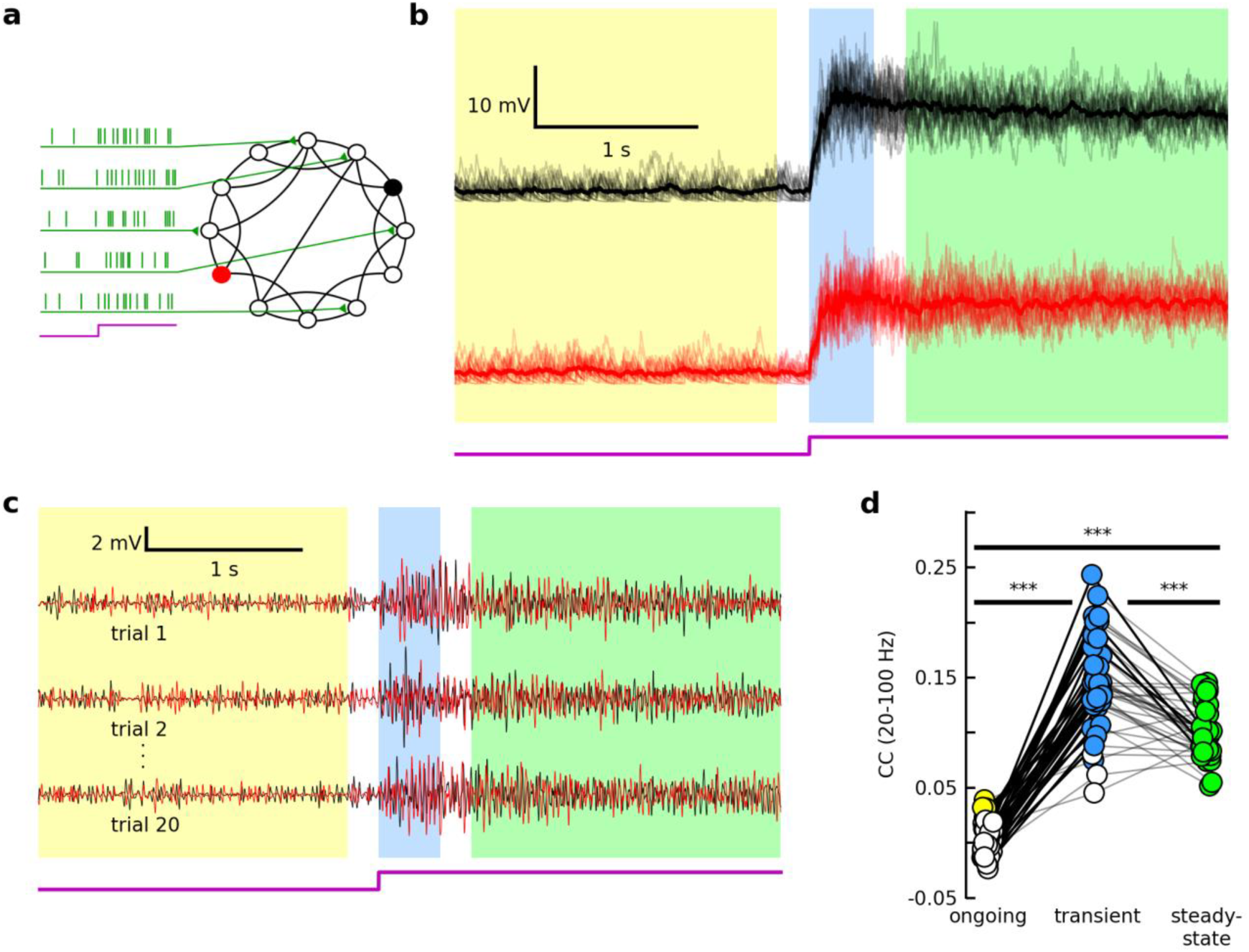
A model network strengthens the “internal mechanism” hypothesis, suggesting crucial roles for network oscillations. (a) The model network was composed of 800 excitatory LIF neurons with small-world connectivity, and 200 randomly-connected inhibitory LIF neurons (not depicted). All excitatory neurons received Poisson external inputs, and the stimulus was modeled as a gradual increase in the external input rate (see Methods). (b – c) Same as in Fig. 3b, c, but two excitatory model neurons randomly-selected from the network (see Methods). (d) Same as in Fig. 3d, but for forty pairs of excitatory neurons, generated from twenty neurons randomly-selected from the network.

In addition, the model network reproduced the experimentally-observed dynamics of correlated variability. The simulated correlated variability increased significantly from ongoing to transient (<CC> = 0.003 ongoing, 0.146 transient, P = 3.6 x 10^−8^ for ongoing – transient comparison) (Fig. 5d), a trend that was consistent with our experimental results (Fig. 3b, c). Mediated by synaptic depression, correlated variability decreased significantly from transient to steady state (<CC> = 0.10 steady-state, P = 1.49 x 10^−6^ for transient – steady-state comparison, P = 3.6 x 10^−8^ for ongoing – steady-state comparison) (Fig. 5d). In contrast, when synaptic depression was absent, correlated variability increased from transient to steady state (**Supplementary Fig. 7a**).

As observed in experiment, this dynamic of correlated variability depended crucially on network activity oscillations. When synaptic time constants for excitation and inhibition were chosen to be identical (see **Methods** and **Supplementary Information 5**), the stimulus instead pushed the network into a regime of asynchronous activity, as evidenced by the population spike-rate and residual membrane potential power spectra (**Supplementary Fig. 6c**). This had little effect on the overall network activity level and gamma power dynamics (**Supplementary Fig. 6d**). Correlated variability, however, was very weakly modulated by the stimulus (**Supplementary Fig. 7b**), in contrast with our experimental results (Fig. 3b, c). This dependence on network oscillations had a synaptic basis: in the synchronous model network, not only were correlations between synaptic conductances stronger than those in the asynchronous network, the lag between excitation and inhibition was larger, allowing for a longer temporal window in which synaptic input correlations could be manifested in pairs of membrane potentials^3,6^ (**Supplementary Fig. 8**). In conclusion, changes in network oscillations, rather than network activity levels alone, determine the changes in correlated variability.

The network oscillation dynamic described here is not the only one capable of reproducing the experimentally-observed dynamic of correlated variability. For example, a similar randomly-connected network generates oscillations that either invade the entire network or are absent, and the CC dynamic can be reproduced by abolishing the oscillation early in the transient window via strong synaptic depression (data not shown). This oscillation dynamic is inconsistent with the experimental results, in which gamma oscillations were strongest during the transient epoch, but persisted in the steady-state (Fig. 3a, 4a, **Supplementary Fig. 4a**). In contrast, the small-world network reproduces the correct network oscillation temporal dynamics (Fig. 5c, **Supplementary Fig. 6a, bottom, 9a**). Furthermore, the spatial arrangement of connections introduced by the small-world architecture adds a spatial dimension to the dynamics of network activity (see **Supplementary Information 5**, **Supplementary Fig. 9a**). Soon after stimulus onset, when synaptic connections are near their default strength, oscillatory spiking is largely coherent across the network of excitatory neurons. However, because of synaptic depression and the distance-dependence of connections, the network dynamically subdivides, and coherent oscillations proceed among spatially separate groups of neurons in the steady-state. This spatiotemporal dynamic predicts a dependence of CC temporal dynamics on distance between neurons (**Supplementary Fig. 9b, c**).

Taken together, these results strengthen the hypothesis that the experimentally-observed dynamics of correlated variability are primarily driven by the dynamics of thalamocortical network oscillations. As such, synaptic time constants and synaptic depression are two intrinsic parameters relevant to correlated variability. Constraining the model to also produce a realistic network oscillation temporal dynamic reveals an additional role for synaptic clustering.

### Discussion

To study how cortical coordination evolves during visual processing, we measured correlated variability between the membrane potentials of pyramidal neuron pairs in turtle visual cortex during ongoing and visually-evoked activity. This approach provides an exceedingly rare and much-needed view into the subthreshold events underlying coordinated spiking activity^6^, and importantly, is uncorrupted by issues associated with spike-based investigations. For example, the process for targeting cells for recording is agnostic to the cells’ spike rates, and this study thus gives voice to the “dark” neurons that overwhelmingly populate the cortex^24^, but are rarely represented in the vast literature on correlated variability. By supplementing our experimental approach with a model network investigation, we further addressed the relative contribution of the thalamocortical network to the dynamics of correlated variability, and identified relevant network variables.

We found that both continuous and brief visual stimulation evoked large, low-frequency membrane potential fluctuations (Fig. 1e), with nested gamma-band (20 – 100 Hz) oscillations (Fig. 2c, 3a), both of which varied significantly from trial to trial (Fig. 2a, b, 3a). This gamma-band variability was significantly correlated across the population of pairs in a given window of activity (Fig. 2b, c). It is reasonable to ask whether this is at all relevant to sensory processing, given that cortical neurons transmit spikes, and not subthreshold fluctuations. Previous work suggests it is. First, the fluctuations themselves are important for interpreting spiking activity in networks of sparse-spiking neurons; subthreshold gamma oscillations define narrow windows in which a given neuron is most likely to fire, and thus determine precise spike timing^25,26^. Second, the across-trial variability of these fluctuations constrains that of the spiking activity^27^; spiking responses can be no more reliable than the corresponding subthreshold activity. Finally, if and when neurons do spike in this visually-evoked “high conductance” state, the spike correlations will be shaped by state-induced changes in response gain, distance from threshold, and subthreshold correlations^6^. In fact, this last relationship may be supralinear^28^, meaning that small changes in subthreshold correlated variability can have major repercussions for supra-threshold coordination.

What determines the strength of correlations in these single-trial deviations for a given pair of neurons? Anatomical connectivity is one obvious candidate, but what are the relative contributions from feedforward and recurrent connections (Fig. 1b)? Recent work has shown that the coupling of the spiking activity of a neuron with that of the population is stimulus-invariant^4,5^, suggesting a crucial role for intracortical connectivity. In agreement with this, we found that gamma band correlated variability for ongoing and evoked activity was significantly related, for both the non-adapted (transient, Fig. 2e) and adapted (steady-state, Fig. 2f) conditions. In addition, although all pairs of neurons were separated by less than 300 microns, and therefore likely received similar sensory inputs^29^, correlated variability was broadly-distributed across the population during a given window of activity (Fig. 3b, c). This variability suggests that the sparse connectivity of the cortex dramatically affects the distribution of pairwise correlations^3^. Indeed, alternate network models with higher connectivity were unable to reproduce this variability (data not shown).

Connectivity was not the only relevant parameter; for most pairs of neurons, and for the population as a whole, gamma band correlated variability increased with visual stimulation, but then relaxed to near ongoing values, despite persistent activity (Fig. 2). That is, for a given network (i.e., an anatomical arrangement of feedforward and recurrent connections), the level of coordination was not static. This was true for responses to both continuous and brief stimulation (Fig. 3b, c) suggesting the dynamic did not simply reflect a change in the statistics of the sensory inputs. Previous experimental work implicates the cortical state: coordination in spiking activity^4,5^, synaptic inputs^8,30^, and membrane potentials^26^ appear to be related to network synchrony. Accordingly, we found ample experimental and computational evidence implicating an evolving network state. Specifically, changes in correlated variability across epochs were related to changes in properties of membrane potential power spectra (Fig. 4b, c), and reflected a phase synchrony dynamic (**Supplementary Fig. 4**) consistent with a network adapting from a “disinhibited” to a “balanced” state^31^. In addition, the model revealed a strong relationship between network spike-rate oscillations and gamma-band correlated variability (**Supplementary Information 5**), and also demonstrated the synaptic basis: synaptic current correlations and the lag between excitation and inhibition are larger for pairs of neurons in a synchronous network than for those in an asynchronous network (**Supplementary Information 5**, **Supplementary Fig. 8**). As such, our experimental results relating correlated variability to network state are in simultaneous agreement with multiple experimental studies of cortical activity across a variety of preparations and spatial scales, and importantly, demonstrate this principle in action at the level of the membrane potential during sensory processing. Moreover, our model results confirm computational predictions relating synaptic current dynamics to membrane potential correlations^3,6^, and extend previous work by demonstrating the effects of synaptic clustering and adaptation.

Gamma band correlated variability decreased from transient to steady-state, yet remained significantly nonzero (Fig. 2d, 3c). While vanishingly-small values would theoretically support greater response fidelity^9^, this realized steady-state value may reflect the level of neuronal interaction necessary for cortical function (e.g., feature binding^32^, effective signal propagation^33^, and general cognitive function^34^). In other words, cortical coordination during sensory processing is maintained at an intermediate level (i.e., less than that during the initial response phase, but larger than zero) that represents the ideal balance between competing cortical needs (Fig. 1c). This balanced state can be maintained across stimulus conditions by intrinsic adaptation mechanisms.

Previous studies have described a stimulus-induced abolition of low-frequency correlated variability in membrane potential^26^ and membrane potential – LFP^14^ pairs in visual cortex. We observed no such decrease (**Supplementary Fig. 3a, b**). This discrepancy may be partially explained by the nature of ongoing activity. In these previous studies, visual stimulation interrupted large-amplitude, low-frequency events that were coherent across electrodes and had random phase relative to stimulus onset. These fluctuations largely remained in residual traces, likely influencing pre-stimulus correlated variability. In contrast, these spontaneous “bursts” of activity occurred relatively infrequently in our experiments (Fig. 1e, 2a, b). To reproduce this quiescent – to – active sensory-evoked dynamic, we limited the inputs to our model neurons to those from external stimulation, and stimulus-triggered, yet internally-generated events. Our model could likely be modified to reproduce the results of these other works by adding a shared background fluctuation^6^. This brute-force implementation would not be very illuminating, however; spontaneous events can be strikingly similar to evoked^18,19,35^, and may therefore represent activity in the same microcircuits, triggered by events usually hidden from the experimenter. Reproducing spontaneous events in a manner that is consistent with their possible sources is beyond the scope of this study.

Here, we have focused on fundamental properties of the cortex (anatomical and emergent) that are likely to strongly influence correlated variability. Future work can more definitively assess the relative contributions to cortical coordination from various intracortical and extracortical sources not addressed here, such as strong inhibitory feedback^36,37^, thalamic adaptation^38,39^, unequal adaptation of excitatory and inhibitory cortical synapses^40^, and the (time-varying) statistics of neuronal activity in the early visual pathway (e.g., correlations across thalamic inputs^41,42^), to name a few. Of particular interest are top-down influences (e.g., attention), which have been shown to impact spike-count correlations in awake, behaving preparations^43,44^. Our results predict that such higher-order inputs may impact correlated variability by influencing the network state. It will be interesting to test this hypothesis, and to determine the synaptic basis of the interaction if confirmed. Further, it is crucial to test for a relationship between subthreshold correlated variability and psychophysical performance, which will require a behavioral assay. As dual whole-cell recordings in awake, behaving preparations (possibly combined with other recording modalities across multiple areas) become increasingly common, future experiments can be designed to address these exciting questions.

### Methods

#### Surgery

All procedures were approved by Washington University’s Institutional Animal Care and Use Committees and conform to the guidelines of the National Institutes of Health on the Care and Use of Laboratory Animals. Sixteen adult red-eared sliders (*Trachemys scripta elegans*, 150-1000 g) were used for this study. Turtles were anesthetized with Propofol (2mg Propofol/kg), then decapitated. Dissection proceeded as described before^45,46^. In brief, immediately after decapitation, the brain was excised from the skull, with right eye intact, and bathed in cold extracellular saline (in mM, 85 NaCl, 2 KCl, 2 MgCl_2_*6H_2_O, 20 Dextrose, 3 CaCl_2_-2H_2_O, 45 NaHCO_3_). The dura was removed from the left cortex and right optic nerve, and the right eye hemisected to expose the retina. The rostral tip of the olfactory bulb was removed, exposing the ventricle that spans the olfactory bulb and cortex. A cut was made along the midline from the rostral end of the remaining olfactory bulb to the caudal end of the cortex. The preparation was then transferred to a perfusing chamber (Warner RC-27LD recording chamber mounted to PM-7D platform), and placed directly on a glass coverslip surrounded by Sylgard. A final cut was made to the cortex (orthogonal to the previous and stopping short of the border between medial and lateral cortex) allowing the cortex to be pinned flat, with ventricular surface exposed. Multiple perfusion lines delivered extracellular saline, adjusted to pH 7.4 at room temperature, to the brain and retina in the recording chamber.

#### Intracellular Recordings

For whole-cell current clamp recordings, patch pipettes (4-8 MΩ) were pulled from borosilicate glass and filled with a standard electrode solution (in mM; 124 KMeSO_4_, 2.3 CaCl_2_-2H_2_O, 1.2 MgCl_2_, 10 HEPES, 5 EGTA) adjusted to pH 7.4 at room temperature. Cells were targeted for patching using a dual interference contrast microscope (Olympus). Simultaneously recorded cells were located less than 300 microns apart, and all cells were located within 300 microns of an extracellular recording electrode. Intracellular activity was collected using an Axoclamp 900A amplifier, digitized by a data acquisition panel (National Instruments PCIe-6321), and recorded using a custom Labview program (National Instruments), sampling at 10 kHz. We excluded cells that did not display stable resting membrane potentials. The visual cortex was targeted as described below.

#### Extracellular Recordings

Extracellular recordings were achieved with tungsten microelectrodes (MicroProbes heat treated tapered tip), with approximately 0.5 MΩ impedance. Electrodes were slowly advanced through tissue under visual guidance using a manipulator (Narishige), while monitoring for activity using custom acquisition software (National Instruments). Extracellular activity was collected using an A-M Systems Model 1800 amplifier, band-pass filtered between 1 Hz and 20,000 Hz, digitized (NI PCIe-6231), and recorded using custom software (National Instruments), sampling at 10 kHz.

#### Identification of Visual Cortex

We used a phenomenological approach to identify the visual cortex, described previously^47^. In general, this region was centered on the anterior lateral cortex, in agreement with voltage-sensitive dye studies^48,49^. Anatomical studies identify this as a region of cortex receiving projections from lateral geniculate nucleus^29^.

#### Visual Stimulation

Whole-field flashes were presented using either a red LED (Kingbright, 640nm), mounted to a manipulator and positioned 1 – 5 cm above the retina, or a projector-lens system (described below). The mean LED light intensity (irradiance) at the retina was 60 W/m^2^. For one turtle, we used these same LEDs in conjunction with 200 micron optical fibers (Edmund Optics) to project sub-field flashes (1 ms – 200 ms) onto the visual streak. Other stimuli were presented using a projector (Aaxa Technologies, P4X Pico Projector), combined with a system of lenses (Edmund Optics) to project images generated by a custom software package directly onto the retina. The mean irradiance at the retina was 1 W/m^2^. This system was used to present brief (100 ms – 250 ms) whole-field and sub-field flashes (red or white), sustained (10 s) gray screen, a naturalistic movie (“catcam”), a motion-enhanced movie (courtesy Jack Gallant), and a phase-shuffled version of the same movie (courtesy Jack Gallant and Woodrow Shew). In all cases, the stimulus was triggered using a custom Labview program (National Instruments).

The preparation was in complete darkness before and after each stimulus presentation. Flashes lasted between 1 ms and 150 ms, with at least 20 s between flashes. Movies lasted either 10 s or 20 s, and were shown at least 12 times, with at least 30 s between the end of one presentation and the beginning of the next.

We presented continuous visual stimuli (movies) while recording from 19 pairs, and brief stimuli (diffuse flashes) while recording from 16 pairs.

#### Signal Processing

In all analyses, only cells with 12 or more visual stimulation trials were included. Raw data traces were down-sampled to 1000 Hz. Because action potentials in turtle cortical pyramidal neurons are relatively wide, spike waveforms still contributed to the band-pass filtered intracellular recordings. To remove these, an algorithm was used to detect spikes, and the membrane potential values in a 20 ms window centered on the maximum of each spike were replaced via interpolation. Finally, the traces were filtered (20 Hz lowpass or 20 – 100 Hz bandpass Butterworth filter).

#### Cross-correlation Analysis

For each single-trial voltage trace, the residual (V_r_ or deviation from the average activity) was found by subtracting the across-trial average time series from the single-trial time series:
*V*_*r*_ = *V* − 〈*V*〉_*trials*_

Residuals were then separated into three epochs: the ongoing epoch (defined to be the two seconds prior to the onset of visual stimulation), the transient epoch (200 to 600 ms after stimulus onset), and the steady-state epoch (800 to 2800 ms after stimulus onset; Fig. 2a). For each pair of simultaneously-recorded cells, the Pearson correlation between residual pairs was then calculated for each epoch and trial. The results were averaged across all trials, resulting in the trial-averaged correlated variability (CC) for each pair and epoch:

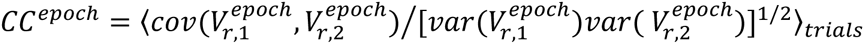

Because the correlated variability of spike counts been shown to depend on the size of the window used for calculations^17^, we repeated the above process for three other sets of choices for epoch window sizes and gaps between epochs (see **Supplementary Information 1**).

The significance of CC for a given pair and epoch was determined by bootstrapping; CC was considered to be significantly nonzero if the average value +/− the 95% confidence level from bootstrapping did not include zero. Similarly, CC for two epochs were considered to be significantly different from one another if the bootstrapping intervals did not overlap.

For the population of pairs, we determined the significance of the population-average CC for a given epoch using the one-sample t-test (that is, by comparing to a zero-mean normal distribution with the same standard deviation). We tested for a significant change in population CC values across two epochs by applying the Wilcoxon signed-rank test to the two sets of CC values.

We also compared CC for responses to brief and continuous visual stimulation. First, pairs were segregated according to the stimulus presented, resulting in 16 brief and 19 extended-stimulus pairs. The two resulting sets of trial-averaged CC values were then compared using the Wilcoxon rank-sum test.

#### Power Analysis

For each trial and cell, we extracted a 5.8 s window of activity (with epoch windows and gaps between epochs as described above, plus 500 ms windows on each end to avoid filtering artifacts in the ongoing and steady-state epochs), and calculated the residual time series as described above. For each residual trace, we performed wavelet analysis in Matlab using software provided by C. Torrence and G. Compo (available at URL: http://paos.colorado.edu/research/wavelets/, ref). This resulted in a power time series for each cell, for multiple frequencies. For each frequency below 100 Hz, we averaged the time series across each epoch to obtain the average power at each frequency for each epoch. We then averaged across trials. For each pair, we also averaged across all frequencies in the gamma range (20 – 100 Hz), and plotted the resulting trial-averaged gamma power (P) in each epoch to inspect for trends across the population (Fig. 2e). We tested for significant changes in a given pair and across the population using the same methods as those described for CC values.

We next inspected for a relationship between changes in CC and changes in gamma power for a given pair. For the ongoing – to – transient and transient – to – steady-state transitions, we calculated the change in trial-averaged gamma power for each neuron

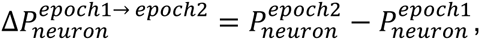

and took the geometric mean of the absolute values:

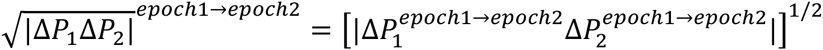

For each pair, we plotted the absolute value of the average change in CC vs. the result, and performed linear regression analysis (Fig. 4a).

For each cell, we also obtained the relative power spectrum (rP) for the transient and steady-state epochs, defined to be the trial-averaged evoked spectrum divided by the trial-averaged ongoing spectrum (Fig. 2d, **Supplementary Fig. 6a**, **c**):

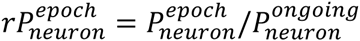

Second, to obtain the “peakiness” of the relative power spectrum within the gamma range for each cell, we divided the maximum value of rP in the gamma range by the average value in the gamma range to obtain the “peak ratio” 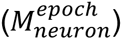 (Fig. 4b, similar to Yu, Ferster 2010):

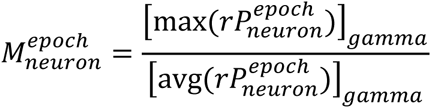

(where 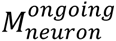 = 1). We then calculated the change in peak ratios across epochs:

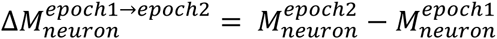

and then the geometric mean of the absolute values for each simultaneously-recorded pair (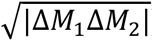). Finally, we inspected for a relationship between 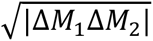 and changes in CC for a given pair (Fig. 4c) as described above for changes in power.

#### Phase Concentration Analysis

For each residual trace in a given trial, we used wavelet analysis (described above) to calculate the phase of each signal as a function of time and frequency. We then averaged over all frequencies in the gamma range (20 – 100 Hz) to obtain the gamma phase time series for each trace (φ_*i*_(t)). For each pair of residuals, we calculated the cosine of the phase difference, and averaged over each epoch to obtain the “phase concentration” (R) for each epoch and trial:

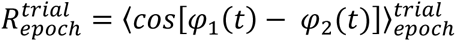

We then averaged over all trials and analyzed the results for pairwise and population trends using the same methods as those described for CC values (**Supplementary Fig. 4b**). We also inspected for a relationship between changes in CC and changes in R for a given pair (**Supplementary Fig. 4c**) as described above for changes in power.

#### Network Models

To investigate the roles of network properties in our experimental results, we implemented a model network of 800 excitatory and 200 inhibitory leaky-integrate-and-fire neurons. Excitatory-excitatory connections had small-world connectivity^41,50^ (with 5% connection probability), and all other connections were random (with 5% excitatory-inhibitory, and 10% inhibitory-excitatory and inhibitory-inhibitory connection probability). Each nonzero entry in the connection weight matrix (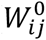) was drawn from a uniform distribution over the interval [0.0, 1.0).

The dynamics of the membrane potential (V) of each neuron evolved according to

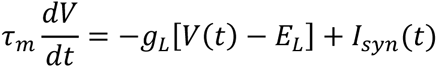

where the membrane time constant τ_M_ = 50 ms (excitatory neurons), 25 ms (inhibitory), and the leak conductance g_L_ = 10 nS (excitatory), 5 (inhibitory). The leak reversal potential EL for each neuron was a random value between −70 and −60 mV, drawn from a Gaussian distribution (to model the variability in resting membrane potentials observed across neurons in the experimental data). The reversal potentials for the synaptic current I_syn_(t) were E_*GABA*_ = −68 mV, and E_*AMPA*_ = 50 mV.

The synaptic current for each synapse type (between presynaptic neurons of type X and postsynaptic neurons of type Y) had three relevant time course parameters: delay (τ*_LX_*, that is, the lag between presynaptic spike time and beginning of conductance waveform), rise time (τ_RYX_), and decay time (τ_DYX_). Synaptic conductances were modeled as products of time-varying gating variables (SYX) and maximum conductances (gYX). Following a presynaptic spike at time 0, the gating variable dynamics were described by

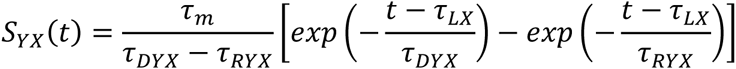

with time constants (in ms) τ_LE_ = 1.5, τ_REE_ = 0.2, τ_DEE_ = 1.0, τ_RIE_ = 0.2, τ_DIE_ = 1.0, τ_LI_ = 1.5, τ_RII_ = 1.5, τ_DII_ = 6.0, τ_REI_ = 1.5, τ_DEI_ = 6.0. Maximum conductance values (in nS) were g_EE_ = 1.0, g_IE_ = 6.0, g_EI_ = 30, g_II_ = 30. In response to a presynaptic spike in neuron *j* at time 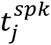, the weight (*W_ij_*) of a synapse connecting neurons *j* and *i* depressed and recovered according to

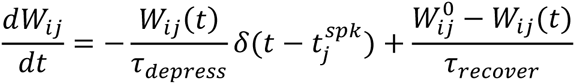

with depression time constant τ_depress_ = 30 ms and recovery time constant τ_recover_ = 1500 ms. Depression and recovery time constants were chosen to give reasonable activity time courses for low-frequency (0 - 20 Hz) membrane potentials.

The spike threshold for each neuron was −40 mV. A neuron reset to −59 mV after spiking, and was refractory for 2 ms (excitatory) and 1 ms (inhibitory).

All excitatory neurons received Poisson external inputs. During “ongoing” activity, the external input rate to each neuron was 25 Hz. The stimulus was modeled as a gradual increase to 500 Hz; the input rate was increased by 95 Hz at stimulus onset, and by an additional 95 Hz every 50 ms for 200 ms. This gradual increase provided more realistic low-frequency membrane potentials than did a single step function stimulus, but did not qualitatively impact the results. The gating variables for external inputs had the same parameters as for excitatory-excitatory connections, and maximum conductances were g_E_ = 4 nS.

Each trial was 5.8 s in duration, with stimulus onset at 2.7 s, and the time step was 0.05 ms. The ongoing epoch was defined to be 2200 ms to 200 ms before stimulus onset, the transient epoch 0 ms to 400 ms after stimulus onset, and the steady-state epoch 600 ms to 2600 ms after stimulus onset. The additional 500 ms at the beginning and end of each trial ensured there were no wavelet filtering artifacts in the ongoing and steady-state epochs.

We then randomly selected 20 excitatory neurons from the entire population of 800 excitatory neurons, and generated 40 V-V pairs from these twenty nodes. Because action potential rates were higher in this model network than in experiment, and because action potentials can affect V-V correlated variability, we substituted “test” neurons for these network neurons before doing the calculation (see **Supplementary Information 6**). Test neurons were identical to network neurons, but all synaptic conductances were multiplied by a factor of 0.5, and spike threshold was raised to −30 mV, which was sufficient to eliminate all action potentials. Thus, a test neuron membrane potential acted as a network sub-sampler, representing the response to re-scaled versions of inputs to the corresponding network neuron. For each pair of test neurons, we then calculated the same parameters as for experimental neuron pairs (CC, R, etc.). We also calculated cross correlations as a function of lag for residual synaptic conductance traces (**Supplementary Figure 8**). This process was identical to that described for the (zero lag) Pearson correlation coefficient described above, but was performed for all lags in +/− 50 ms.

For each trial, we calculated the instantaneous spike rate of all excitatory neurons in the network. We then calculated the power spectra of the transient and steady state spike rates using wavelet filtering, and averaged over all trials (**Supplementary Figure 6**), as described above.

To investigate the dependence on distance between neurons, we repeated the above analysis for this model using two alternate sets of randomly-selected neurons. First, we randomly selected 20 neurons from a group of 100 neighboring neurons. Second, we used a smaller group of 20 neighboring neurons (see **Supplementary Information 5**, and **Supplementary Figure 9**).

In addition, we implemented two alternate model versions (see **Supplementary Information 5**, and **Supplementary Figures 6, 7, 8**). In one, we eliminated synaptic adaptation. In the other, we maintained synaptic adaptation, but tuned synaptic time courses to give an asynchronous transient epoch (τ_LX_ = 1.5 ms, τ_RYX_ = 0.2 ms, τ_DYX_ = 1.0 ms for all X and Y). (In the absence of adaptation, this version was also asynchronous in the steady-state.)

## Acknowledgments

We thank Woodrow Shew for assistance with the design of the visual stimulus. We thank Thomas Crockett for contributing data. This research was supported by a Whitehall Foundation grant #20121221 (R.W.) and a NSF CRCNS grant #1308159 (R.W.).

## Author Contributions

N.W., M.H., and R.W. conceived the study and designed the experiments. N.W. performed the experiments. M.H. performed the model simulations. N.W. analyzed the data. N.W., M.H., and R.W. wrote the paper.

